# Increased glycolysis is an early outcome of palmitate-mediated lipotoxicity

**DOI:** 10.1101/2020.06.10.144808

**Authors:** Pâmela Kakimoto, Antonio Zorzano, Alicia J. Kowaltowski

## Abstract

Palmitic acid is the most abundant saturated fatty acid in human serum. In cell culture systems, palmitate overload is considered a toxic stimulus, and promotes lipid accumulation, insulin resistance, endoplasmic reticulum stress, oxidative stress, as well as cell death. An increased supply of fatty acids has also been shown to change the predominant form of the mitochondrial network, although the metabolic effects of this change are still unclear. Here, we aimed to uncover the early bioenergetic outcomes of lipotoxicity. We incubated hepatic PLC/PRF/5 cells with palmitate conjugated to BSA and followed real-time oxygen consumption and extracellular acidification for 6 hours. Palmitate increased glycolysis as soon as 1 hour after the stimulus, while oxygen consumption was not disturbed, despite overt mitochondrial fragmentation and cellular reductive imbalance. Palmitate only induced mitochondrial fragmentation if glucose and glutamine were available, while glycolytic enhancement did not require glutamine, showing it is not dependent on morphological changes. NAD(P)H levels were significantly abrogated in palmitate-treated cells. Knockdown of the mitochondrial NAD(P) transhydrogenase or addition of the mitochondrial oxidant-generator menadione in control cells modulated ATP production from glycolysis. Indeed, using selective inhibitors, we found that the production of superoxide/hydrogen peroxide at the I_Q_ site of electron transport chain complex I is associated with the metabolic rewiring promoted by palmitate, while not changing mitochondrial oxygen consumption. In conclusion, we demonstrate that increased glycolytic flux linked to mitochondrially-generated redox imbalance is an early bioenergetic result of palmitate overload and lipotoxicity.

## 1. Introduction

High concentrations of nonesterified fatty acids in the plasma are a common feature in the metabolic syndrome, and reflect an incapacity to suppress lipolysis through insulin signaling or inadequate adipose tissue ability to store energetic surpluses (Feng et al., 2017; Hardy et al., 2016; Tchernof and Despres, 2013). Palmitic acid, a sixteen-carbon saturated fatty acid (SFA), is the most abundant saturated fatty acid in human blood (Feng et al., 2017; Quehenberger et al., 2010; Tomita et al., 2011). In excess, palmitate causes lipotoxicity, a disturbance that compromises cell or tissue function, promoted by lipid overload and/or lipid accumulation. In cell culture systems, palmitate overload can promote lipid accumulation, insulin resistance, endoplasmic reticulum (ER) and oxidative stress, as well as cell death (Alsabeeh et al., 2018; Ertunc and Hotamisligil, 2016; Ly et al., 2017). In an unbiased and systematic approach, Piccolis et al. (2019), demonstrated that the most upregulated stress responses after 24 h of palmitate overload are apoptosis, autophagy, and endoplasmic reticulum (ER) stress. The accumulation of glycerolipids, mainly in the di-saturated form, is probably the trigger for toxicity (Masuda et al., 2015; Zhu et al., 2019), but the metabolic pathways sustaining changes induced by palmitate are still not clear.

Mitochondrial function is often suggested as a target for lipotoxicity, since these organelles are central in metabolic control and lipid homeostasis. Mitochondria are dynamic organelles that can quickly adapt to their surroundings with profound changes in function and form (Chan, 2020; Sebastián et al., 2017). In addition to their roles related to ATP production and cell death, mitochondria act as important cell hubs regulating calcium signaling, supplying intermediates for lipid synthesis, and modulating the production and removal of oxidants such as superoxide radical anions. Concomitantly, they sense nutrient availability and cooperate with metabolism-regulating pathways, including responses to insulin and its downstream effectors (del Campo et al., 2014; Cheng et al., 2010; Gordaliza-Alaguero et al., 2019; Wai and Langer, 2016; Zorzano et al., 2015).

Nutrient oversupply predominantly promotes mitochondrial fragmentation, while nutrient starvation is associated with elongation, and a more continuous network (Liesa and Shirihai, 2013; Wai and Langer, 2016; Zorzano et al., 2015). The bioenergetic output is that shortened mitochondria are usually thought to be less efficient in producing ATP than elongated ones (Schrepfer and Scorrano, 2016). Increased supply of fatty acids is an event in which the predominant form of the mitochondrial network is changed. Saturated fatty acid overload can promote mitochondrial fragmentation (Jheng et al., 2012; Kulkarni et al., 2016; Molina et al., 2009) or fusion (Jheng et al., 2012; Senyilmaz et al., 2015), depending on the acyl chain length and if unsaturated fatty acids are also present.

While most studies evaluated later timepoints of palmitate overload (> 12 hours), here we describe the early mitochondrial morphological and bioenergetic outcomes of lipotoxicity. Using real time measurements, we identified that mitochondrial oxygen consumption is not disturbed by palmitate overload over 6 hours, despite high mitochondrial fragmentation and cellular reductive imbalance. Interestingly, palmitate increased ATP synthesis through glycolysis after 1 hour, in a manner unrelated to the changes in morphology. We also found that the production of mitochondrial superoxide radicals/hydrogen peroxide may be associated with this metabolic rewiring promoted by palmitate.

## 2. Materials and methods

### 2.1. Cell cultures

#### 2.1.1. PLC/PRF/5 cells

Human hepatoma PLC/PRF/5 cells, or “Alexander cells”, were maintained in Dulbecco’s Modified Eagle Medium (DMEM) containing 5.5 mM glucose, 4 mM glutamine, 1 mM pyruvate, 25 mM Hepes, 1% penicillin/streptomycin, and 10% fetal bovine serum, in a humidified, 5% CO_2_, 37°C incubator.

#### 2.1.2 Nicotinamide nucleotide transhydrogenase knockdown

Nicotinamide nucleotide transhydrogenase (NNT) knockdown was performed on PLC/PRF/5 cells as described in Muñoz and Zorzano, 2015, with some modifications. Briefly, HEK293T cells were seeded on 10 cm plates in DMEM containing 25 mM glucose, 4 mM glutamine, 1 mM pyruvate, 25 mM Hepes, 10% fetal bovine serum, and kept in a humidified, 5% CO_2_, 37°C incubator. Twenty-four hours later, cells were transfected for lentiviral production in media containing 5 μg of each lentiviral packing plasmid (pMD2.G and psPAX2, Addgene: #12259, #12260, respectively), 10 μg of the scramble pLKO.1 or the NNT constructs (named here 1 and 2, respectively, from Mission® shRNA: TRCN0000028512, TRCN0000028541), and PEI Max, then incubated for an additional 24 hours at 37°C. The following day, media was replaced and cells were incubated at 33°C for viral production. Twenty-four hours later, media was collected, filtered in a 0.45 μm filter, and mixed with 2.5 μg.mL^-1^ of Polybrene for PLC/PRF/5 infection. A second infection was performed the following day. Puromycin-resistant cells were selected in complete DMEM supplemented with 2.5 μg.mL^-1^ puromycin. After selection, cells were expanded and maintained in complete DMEM supplemented with 2.5 μg.mL^-1^ puromycin.

### 2.2. Fatty acid conjugation to bovine serum albumin

Sodium palmitate was solubilized in MilliQ water by heating (70°C) and conjugated to fatty acid-free bovine serum albumin (FFA BSA) at 37°C. A stock solution (4 mM fatty acid in 5% albumin) was filtered (0.22 μm) and frozen under sterile conditions at −20°C for later use.

### 2.3. Cell treatments: fatty acids

Cells were plated in maintenance DMEM. After 24 hours, the cells were PBS-washed and the medium was replaced by DMEM in the presence or absence of 5.5 mM glucose and 4 mM glutamine, supplemented with sodium palmitate, oleate, or stearate conjugated to FFA BSA to final a concentration of 200 μM in 0.25% FFA BSA. Control groups were incubated in 0.25% FFA BSA.

### 2.4. SDS Page and western blots

After incubation with fatty acids, cells were lysed in RIPA buffer and centrifuged (16000 g, 30 minutes). The supernatant was collected and protein quantification was performed using Pierce’s BCA reagent and a BSA standard curve. Proteins were diluted in Laemli buffer containing 5% β-mercaptoethanol and heated for 5 minutes for denaturation. Proteins were separated by electrophoresis in a denaturing polyacrylamide gel and transferred to a polyvinylidene difluoride (PVDF) membrane. After blocking with 5% defatted milk, membranes were incubated (overnight, 4°C) with primary antibodies (anti-NNT cat. Abcam: ab110352; β-actin cat. Abcam ab8226). Membranes were then incubated with secondary antibodies conjugated to fluorescent (Licor®) anti-mouse.

### 2.5. Confocal microscopy

Mitotracker Deep Red (50 nM, ThermoScientific^®^) or methyl tetramethylrhodamine ester (TMRM 100 nM, ThermoScientific®) and Bodipy 493/503 (0.2 μg.mL^-1^, ThermoScientific®) were added to the cells (plated on 4-chamber CELLView, Greiner BioOne®) and incubated 30 minutes at 37°C and 5% CO_2_. Immediately after, live cell images were acquired using a Zeiss LSM 880 Elyra Ayriscan confocal microscope, (temperature, humidity and CO_2_ control). The acquisition was made in 35 positions on the Z axis, in order to enable reconstruction in 3 dimensions, or in 7 positions on the Z axis and 9 XY blocks, in order to quantify the population.

#### 2.5.1. Image analysis

The mitochondrial network was reconstructed in 3D with the aid of MitoKondrY macro for ImageJ, developed by Dr. Sebastien Tosi (IRB Barcelona). Briefly, the macro allows quantification in pixels of the volume and the length of the mitochondrial network. From that, objects that do not form branches (not contained in the network) are identified, filtered as “individuals” for further quantification of their volume, surface area, axis length, sphericity and aspect ratio (major axis/minor axis). Mitochondrial population evaluation took place after blind identification and sample randomization. Previous definitions were followed: predominantly fragmented - classified as “short” - or predominantly elongated - called “long”. The cells called “intermediate” present a network with some proportion of both the shortened and elongated phenotypes. Cells were classified and counted for later identification (Muñoz and Zorzano, 2015). The mitochondrial fusion factor (FF) is defined as % intermediate cells + 2*% elongated cells (Senyilmaz-Tiebe et al., 2018).

### 2.6. Lactate production

After cell incubation on P24 plates, media were collected and the cells were PBS-washed and frozen. Secreted lactate content was measured in the media using a colorimetric assay, following the manufacturer’s instructions (LabTest®, Brazil). Values were normalized by total cell protein content.

### 2.7. Triglyceride content

After cell incubation on 10-cm plates, cells were PBS-washed and scraped in 1 mL PBS. Eight hundred microliters were centrifuged and 200 μL were kept for protein quantification. After centrifugation (5 min, 300 g), lipids were extracted by Folch’s method (adapted from Folch et al., 1957; Wang et al., 2017). Briefly, the pellets were resuspended in 1 ml chloroform:methanol (2:1) and vigorously agitated for 1 hour. Two-hundred microliters of MilliQ water were added to the tube and vortexed. The organic phase was collected to a new tube and dried overnight in the fume hood. The pellet was resuspended in 200 μL chloroform and 2% Triton-X 100 and re-dried. The remaining pellet was resuspended in 50 μL MilliQ water and the triglyceride content was measured by a colorimetric assay, following the manufacturer’s instructions (LabTest^®^, Brazil). Values were normalized to the total cell protein content.

### 2.8. Metabolic analysis: oxygen consumption, extracellular acidification rates, and ATP production

#### 2.8.1. Palmitate overload: oxygen consumption and extracellular acidification rate time courses

Twenty-four hours after plating on XFe24 Seahorse plates (Agilent^®^), cells were PBS-washed and incubated in DMEM containing 5.5 mM glucose, 4 mM glutamine, 1 mM pyruvate, 1% penicillin/streptomycin, and 5 mM Hepes. Media did not contain bicarbonate nor FBS. Cells were kept for 30 minutes in a humidified, 37°C incubator. After basal oxygen consumption and extracellular acidification rates (OCR and ECAR, respectively) were measured, sodium palmitate conjugated to FFA BSA was injected into the wells to final concentrations of 200 μM/0.25%. Control groups received 0.25% FFA BSA. Measurements were taken, after the injection, and at every hour, up to 6 hours. Cell-free wells were incubated with BSA or palmitate/BSA for background correction.

#### 2.8.2. Palmitate overload: glycolysis modulation with 2-deoxyglucose

Twenty-four hours after plating on XFe24 Seahorse plates (Agilent^®^), cells were PBS-washed and incubated in DMEM containing 5.5 mM glucose, 4 mM glutamine, 1 mM pyruvate, 1% penicillin/streptomycin, 200 μM/0.25% palmitate/FFA BSA, 5 mM Hepes. Control groups received 0.25% FFA BSA. Media did not contain bicarbonate nor FBS. Cells were kept for 6 hours in a humidified 37°C incubator. OCR and ECAR were measured under basal conditions and following the addition of 20 mM 2-deoxyglucose, 1 μM oligomycin (pre-titrated), 1 μM rotenone and 1 μM antimycin A. ATP production rates (glycolytic, mitochondrial, and total) were calculated from the ECAR, OCR, and proton exchange rate (PER = ECAR*BF*Vol_microchamber_*K_vol_), following the manufacturer’s instructions. Briefly, mitochondrial-derived ATP is (OCR_basal_ – OCR_oligomycin_)*P/O. Glycolytic ATP is PER_Total_ – (OCR_basal_ – OCR_Rot/AA_)*CCF. Total ATP production is the sum of mitochondrial and glycolytic ATP. We used the manufacturer’s standard values for K_vol_, P/O (phosphate/oxygen ratio) and CCF (CO_2_ contribution factor). BF (buffer factor) was previously measured as 3.13 mM · pH^-1^.

#### 2.8.3. Palmitate overload: glucose and/or glutamine deprivation

Twenty-four hours after plating on XFe24 Seahorse plates (Agilent^®^), cells were PBS-washed and incubated in DMEM containing, 1 mM pyruvate, 1% penicillin/streptomycin, 200 μM/0.25% palmitate/FFA BSA, and 5 mM Hepes. Eight groups were assigned for nutrient deprivation experiments (with or without palmitate): i) N/A: without glucose and glutamine, ii) Glc+Gln: added glucose and glutamine, iii) Glc: added only glucose, iv) added only glutamine. Concentration for glucose and glutamine were 5.5 mM and 4 mM, respectively. Control groups received 0.25% FFA BSA. Media did not contain bicarbonate nor FBS. Cells were kept for 6 hours in a humidified 37°C incubator. Oxygen consumption and extracellular acidification rates were measured under basal conditions, and with 1 μM oligomycin followed by 1 μM rotenone and 1 μM antimycin A. ATP production rates were calculated as described in the manufacturer’s instructions. The buffer factors were previously measured for each buffer (in mM · pH^-1^): Glc+Gln = 3.13, Glc = 3.34, Gln = 3.13, Noadditions = 3.6.

#### 2.8.4. Palmitate overload: oxidant production modulation

Twenty-four hours after plating on XFe24 Seahorse plates (Agilent^®^), cells were PBS-washed and incubated in DMEM containing 5.5 mM glucose, 4 mM glutamine, 1 mM pyruvate, 1% penicillin/streptomycin, 200 μM/0.25% palmitate/FFA BSA, 5 mM Hepes, and 10 μM S1QEL 1.1, S3QEL 2, or menadione. Control groups received 0.25% FFA BSA and/or 0.1% DMSO as vehicle. Media did not contain bicarbonate nor FBS. Cells were kept for 6 hours in a humidified 37°C incubator. Oxygen consumption and extracellular acidification rates were measured under basal conditions, and after the addition of 1 μM oligomycin, followed by 1 μM rotenone and 1 μM antimycin A. ATP production rates were calculated as described in the manufacturer’s instructions.

#### 2.8.5. Protein normalization

After every run, the media was removed and the cells were frozen. Post-thawing, cells were disrupted by RIPA buffer and vigorous agitation. Protein quantification was performed using Pierce’s BCA reagent and a BSA standard curve.

### 2.9. NAD(P)H fluorescence

After 6 h in 200 μM/0.25% palmitate/FFA BSA or 0.25% FFA BSA, cells plated on P100 plates were PBS-washed and tripsinized. Cells were resuspended in DMEM with 5.5. mM glucose, 4 mM glutamine, 1 mM pyruvate and 5 mM Hepes, and counted. Half a million cells per milliliter were incubated in DMEM containing 200 μM/0.25% palmitate. NAD(P)H fluorescence (excitation 366 nm, emission 450 nm) was followed in a F4500 Hitachi^®^ fluorimeter with magnetic stirring. Cells were challenged with two sequential additions of t-butyl hydroperoxide (0.5 mM) every 5 minutes. After 3 minutes, oligomycin (1 μM), carbonyl cyanide 3-chlorophenylhydrazone (CCCP, 1 μM – to promote maximal oxidation), and rotenone (1 μM) plus antimycin A (1 μM), to promote maximal reduction, were sequentially added.

### 2.10. Statistics

Analysis was conducted using Microsoft Excel, RStudio, GraphPad Prism or JASP. Outliers were removed by Grubb’s test (alpha = 0.05). If the samples were normally distributed (p > 0.05, Shapiro-Wilk’s test) and the variances were equal (p > 0.05 Levene’s test), the group’s average was compared by Student’s t test or two-way Analysis of Variance (ANOVA) followed by Tuke’s posttest. Levels of significance were considered at p < 0.05.

### 2.11. Materials

Culture media and supplements were purchased from Thermo Scientific, except DMEM without glucose (cat. D5030), that was from Sigma-Aldrich. Oligomycin, CCCP, rotenone, and antimycin A were from Santa Cruz Biotechnology. All the other reagents were purchased from Sigma-Aldrich, unless stated differently.

## 3. Results

### 3.1. Palmitate promotes mitochondrial fragmentation and triglyceride accumulation

Palmitate overload promotes mitochondrial fragmentation in many cell types (Jheng et al., 2012; Kulkarni et al., 2016; Molina et al., 2009), but the relationship between these morphological changes and early functional bioenergetic consequences has not been established. We observed that exposure to 0.2 mM palmitate for 6 h led to overt mitochondrial fragmentation in human hepatoma PLC/PRF/5 cells (Fig 1A shows a typical image, quantified in Fig 1B). Prolonged stimuli of 24 hours increased the fraction of short mitochondria (Fig 1C-D). These morphological changes were confirmed by the quantified decrease of mitochondrial fusion factor (Fig 1E) and by the automated quantification of the mitochondrial population in which individual sphericity is increased (Fig 1F). Importantly, no changes were detected in total mitochondrial network volume (Fig 1G). In parallel, we verified that 6 hours of palmitate treatment were enough to induce significant triglyceride accumulation (Fig 1H). These results confirm the ability of palmitate overload to replicate mitochondrial fragmentation and lipid accumulation in PLC cells, validating it as a lipotoxicity model, with strong results detected as soon as 6 hours of incubation.

**Figure 1:**
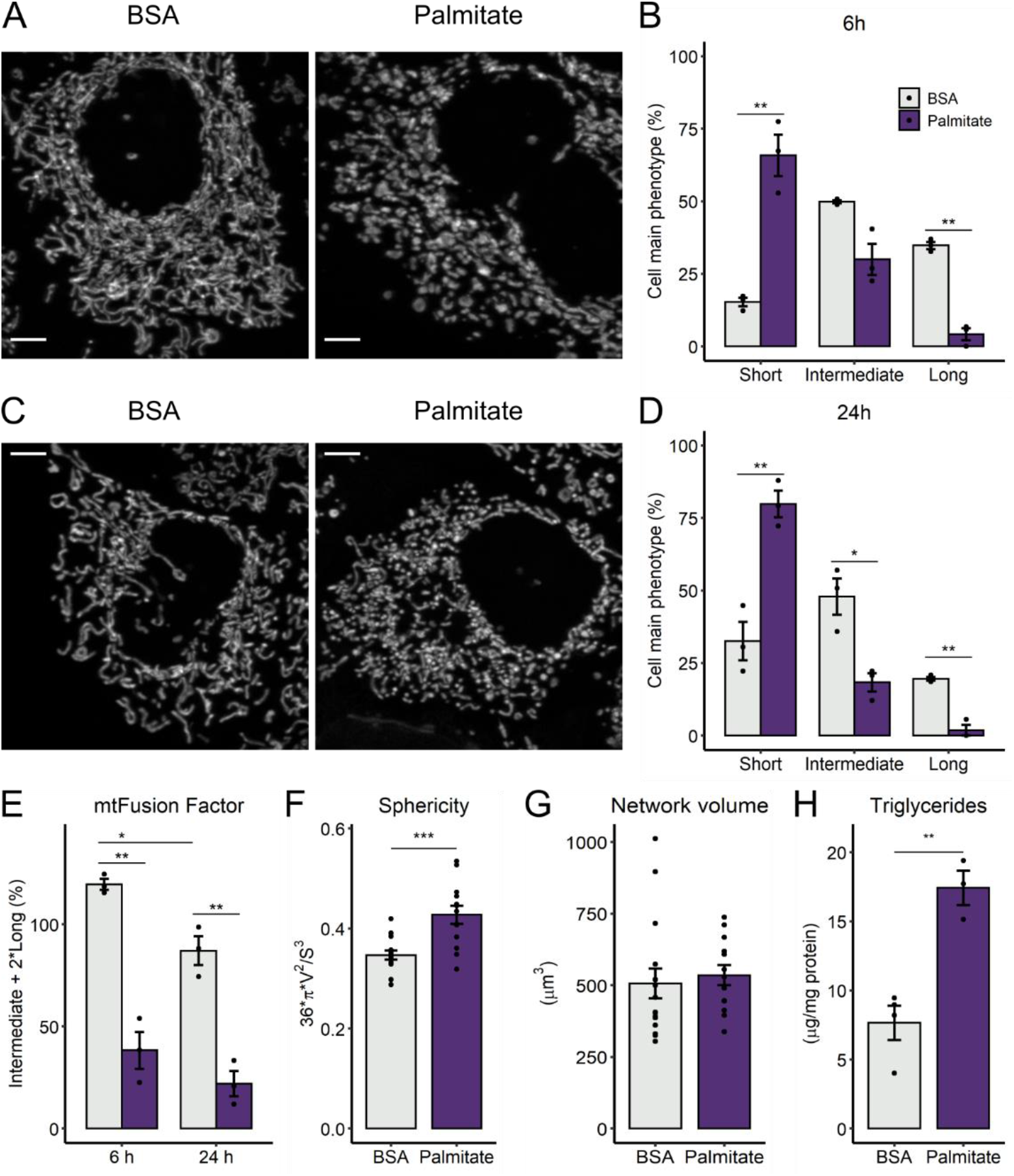
Palmitate overload promotes mitochondrial fragmentation and triglyceride accumulation. PLC/PRF/5 cells were incubated with palmitate/BSA (0.2 mM/ 0.25%) for 6 h or 24 h. Live mitochondrial morphology was evaluated by confocal microscopy in cells loaded with 0.4 μM TMRM at 6 h (A and B) or 24 h (C and D). Morphological parameters were measured as described in the Material and Methods section, in cell populations (B, D, E) or individual cells (F-G). Triglycerides were extracted and quantified after 6 h of palmitate incubation (H). * = p < 0.05, ** = p < 0.01, *** = p < 0.001 t-test (Panels B, D, F-H) or two-way ANOVA followed by Tukey’s post-test (E). Data are mean + SD. Filled circles represent independent experiments. In F-G, filled circles represent individual cells obtained from 3 independent experiments. Scale bar = 5 μm.

### 3.2. Palmitate induces glycolysis

Changes in the predominant shape of the mitochondrial network as a response to their microenvironment have been associated with modifications of mitochondrial bioenergetic capabilities, i.e., ATP production (Chan, 2020; Schrepfer and Scorrano, 2016). We followed real-time respiration of intact cells just after palmitate treatment, and up to 6 hours (Fig 2A). Surprisingly, oxygen consumption rates (OCRs) of palmitate-treated cells were not significantly affected over time compared to the BSA controls (Fig 2A). However, palmitate promoted a significant increase in extracellular acidification rates (ECARs; Fig 2B).

**Figure 2:**
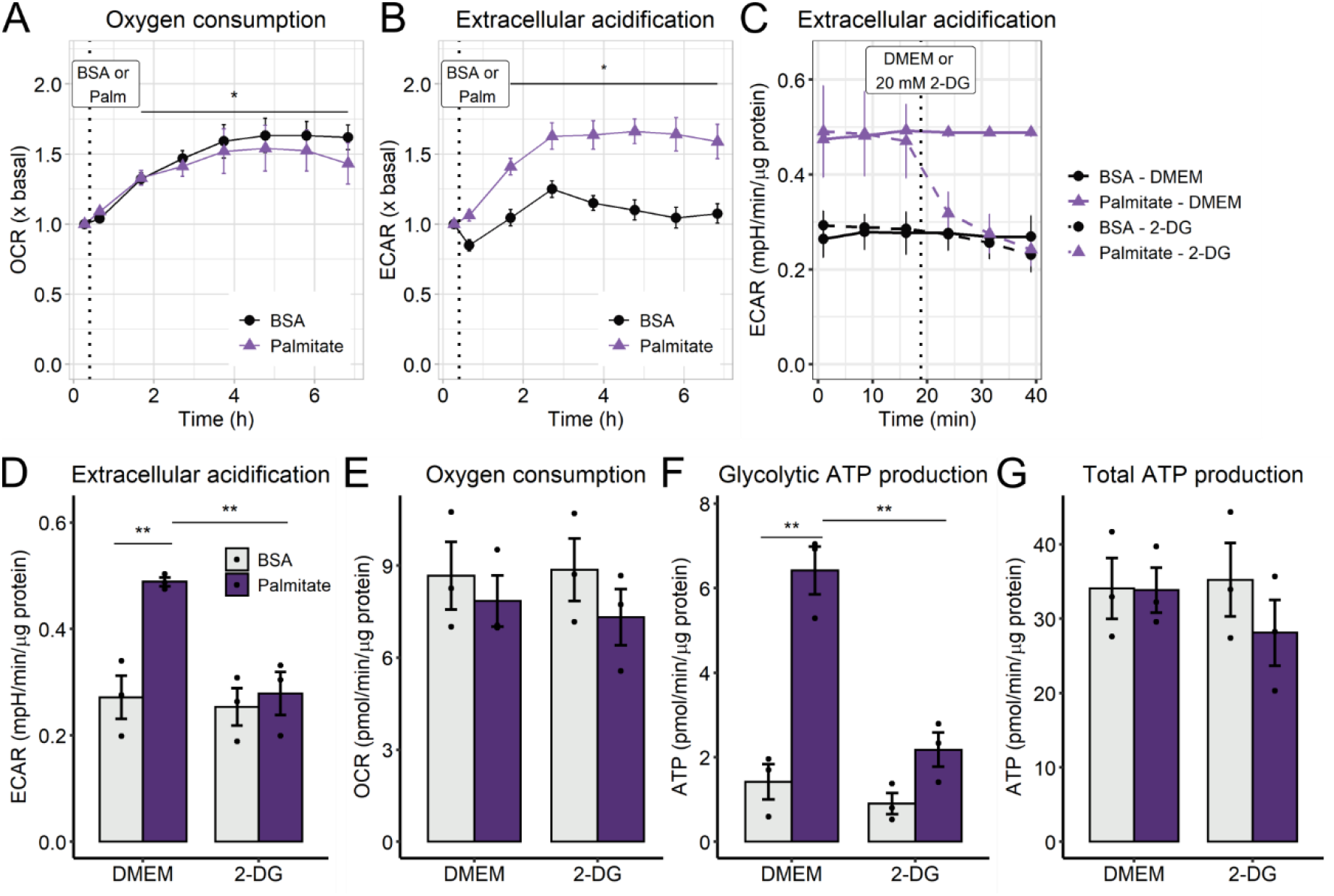
Palmitate overload increases glycolytic flux. Real-time OCRs and ECARs were measured in a Seahorse XFe24 analyzer over 6 h after 0.2 mM palmitate injection, indicated by the dotted line (A-B). ECARs and OCRs were measured after 6 h palmitate incubation and modulated by 2-deoxyglucose (C-E). Glycolytic ATP and total ATP production were calculated as described in the Methods section (F-G). In A and B, * = p < 0.05 for time effect, two-way ANOVA repeated measurements. In D-G, ** = p < 0.01 two-way ANOVA followed by Tukey’s post-test. OCR and ECAR plots are means + SEM. Bar plots are means + SD. Filled circles represent independent experiments.

Differences in ECARs can be related to lactate production, but also to changes in tricarboxylic acid cycle flux (due to CO_2_), ATP turnover, or other acid-generating catabolic processes (Mookerjee et al., 2015). To evaluate the source of protons, we monitored the ECARs in the presence of 2-deoxyglucose, an inhibitor of hexokinases. In palmitate-treated cells, 2-deoxyglucose blunted the ECAR increase (Fig 2C-D) without changing OCR (Fig 2E), but with an associated decrease in ATP production from glycolysis (Fig 2F). Importantly, palmitate treatment very strongly increments the contribution of glycolysis in ATP production (Fig 2F), while maintaining total ATP production equal (Fig 2G). Additionally, palmitate promoted a non-significant tendency toward an increase in lactate production (Suppl Fig 1). This indicates that an early metabolic effect of palmitate treatment is to modulate the sources of ATP production, surprisingly increasing glycolytic ATP generation. This change involves metabolic fluxes only, since the protein content of GLUT1 and GAPDH was unchanged (data not shown). From these results, we can conclude that increased mitochondrial fragmentation precedes any disruptions in mitochondrial oxygen consumption and overall ATP production upon palmitate exposure. Glycolytic flux is increased as soon as 1 h of palmitate treatment, meaning this metabolic switch is an early outcome of palmitate-induced lipotoxicity.

### 3.3. Mitochondrial fragmentation and fuel dependency switch depend on substrate availability

We found it quite surprising that a fatty acid would induce such a significant increase in glycolytic ATP production, and decided to further characterize this effect by examining the role of different substrates present in culture media in this metabolic shift (Fig 3). Cells were treated with palmitate under glucose and/or glutamine deprivation for 6 hours to assess which main media nutrients could be related to metabolic remodeling. Mitochondrial fragmentation was promoted only in full media, i.e., DMEM containing both glucose (Glc) and glutamine (Gln), (Fig 3A-B).Mitochondrial fragmentation was not necessary for the activation of ATP synthesis from glycolysis, as this occurred both in the presence of glucose and glutamine, and with glucose alone (Fig 3C). The presence of palmitate did not change total ATP production in any group, but glutamine deprivation (no additions and Glc groups) reduced it significantly (Fig 3D). Following glutamine deprivation, there was a significant decrease in mitochondrial OCRs (basal, Suppl Fig 3), demonstrating the important role of this amino acid, which feeds into the TCA cycle, to sustain mitochondrial ATP production when glucose is absent (Gln group). From these results, we can conclude that PLC cells have high flexibility for nutrient utilization. Additionally, palmitate does not impede ATP production, but promotes a shift towards glycolysis only if glucose is available.

**Figure 3:**
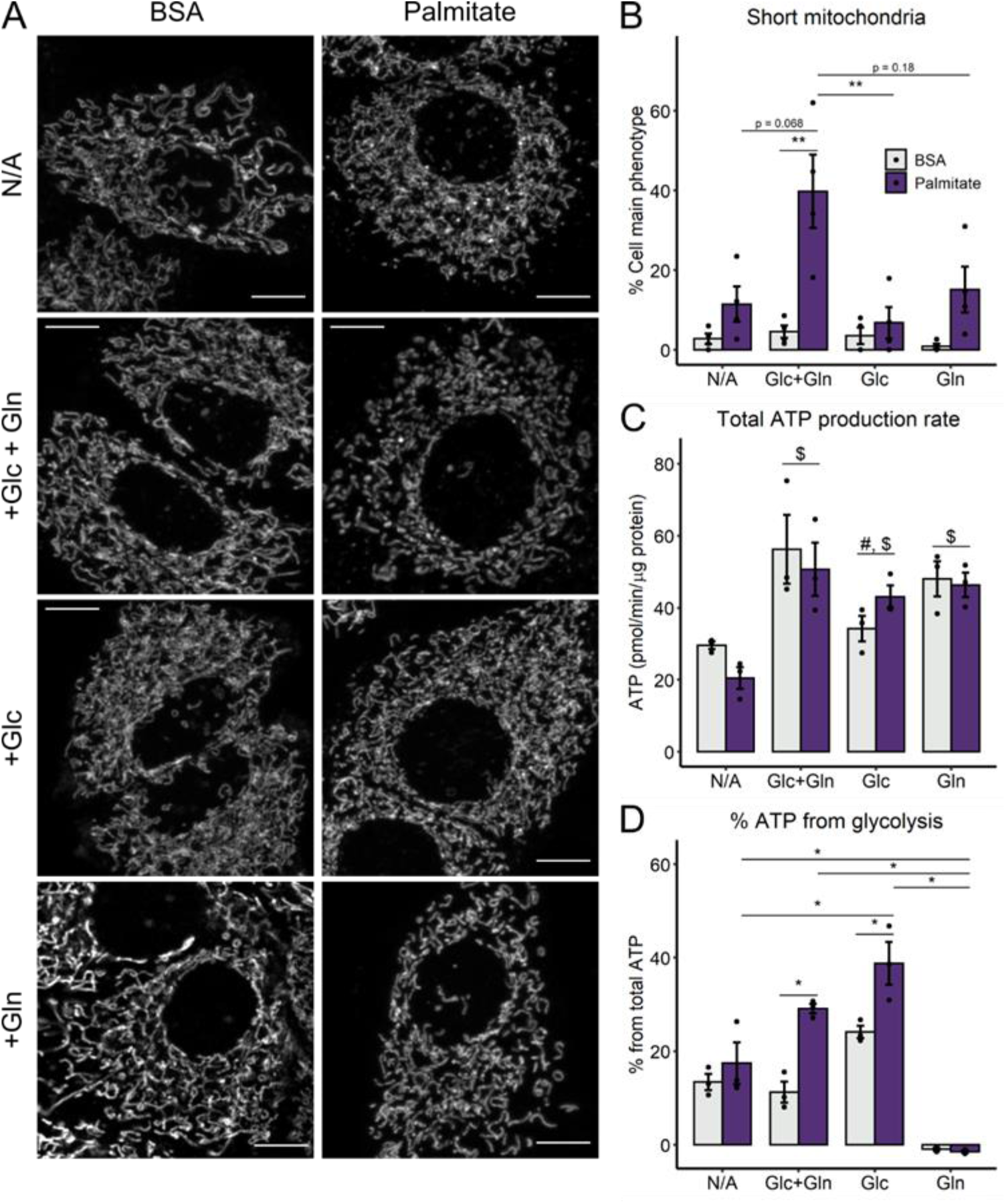
Palmitate-induced metabolic shift requires glucose, but not mitochondrial network fragmentation. PLC/PRF/5 cells were incubated with palmitate/BSA (0.2 mM/ 0.25%) for 6 h in media without glucose nor glutamine (N/A), or in the presence of glucose and glutamine (Glc+Gln), only glucose (Glc), or only glutamine (Gln). Live mitochondrial morphology was evaluated after 50 nM Mitotracker Deep Red loading by confocal microscopy at 6 h (A). Mitochondrial network morphology was classified as described in the Methods section (B). ATP production was calculated as described in the Methods section (C-D). * = p < 0.05 or ** = p < 0.01; $ p < 0.05 vs. NA; # = p < 0.05 vs. Glc+Gln. Two-way ANOVA followed by Tukey’s post-test. Bar plots are means + SD. Filled circles represent independent experiments. Scale bar = 10 μm.

### 3.4. NAD(P) redox state changes in palmitate-induced metabolic plasticity

Lipid overload is often associated with oxidative stress. The ability of a cell to handle oxidants is related to its NADPH pool, which is essential to maintain reduced glutathione and thioredoxin, involved in the main hydrogen peroxide-removing systems (Winterbourn, 2017). We measured NAD(P)H in cells by autofluorescence, using modulation by the electron transport chain inhibitors oligomycin or rotenone plus antimycin. The uncoupler CCCP was used to maximize reduction and oxidation, and thus estimate the total as well as oxidized and reduced pools (as described in the methods section). Surprisingly, by reducing the total pool of NAD(P)^+^ (through the inhibition of electron transport chain complexes I and III, simultaneously), we observed that palmitate treatment promotes a large increase in total NAD(P) (Suppl Fig 4). Furthermore, under basal conditions, NAD(P)H content is lower in palmitate versus BSA treated cells (Fig 4A). We should note, however, that this assay cannot separate the different cell compartments, as mitochondrial modulation will also change NAD(P)H redox state in the cytosol, since the pools are connected by the reversible malate-aspartate shuttle (Xiao and Loscalzo, 2019). However, our result conclusively indicates that palmitate leads to redox balance changes in PLC cells.

**Figure 4:**
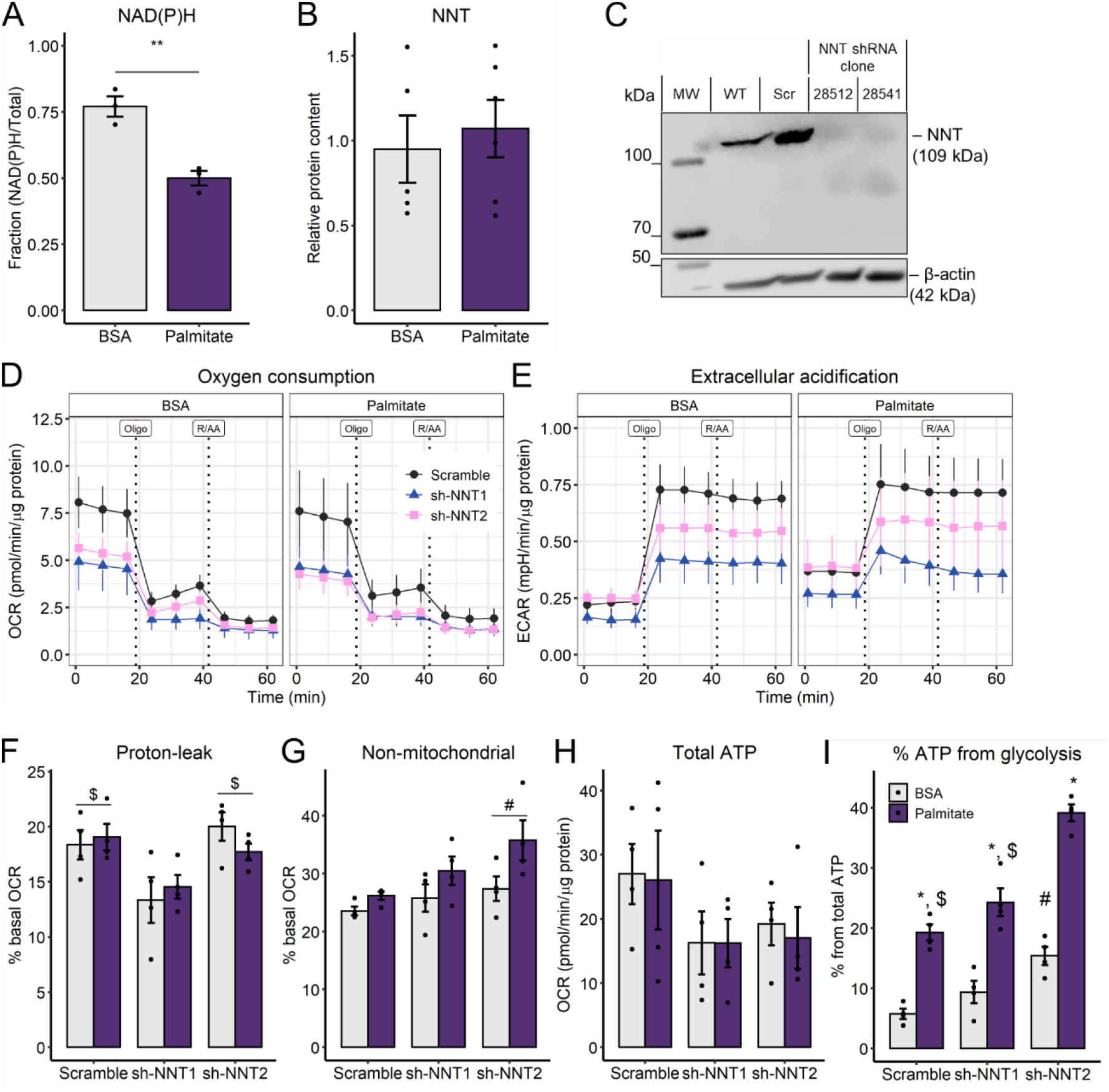
Nicotinamide nucleotide transhydrogenase knockdown exacerbates fuel switch. PLC/PRF/5 cells were incubated with palmitate/BSA (0.2 mM/ 0.25%) for 6 h. Intact cell NAD(P)H content (A), NNT protein content for WT cells (B), or puromycin-selected PLC cells for scramble or NNT knockdown (C) are shown. Real-time OCRs and ECARs were measured in a Seahorse XFe24 analyzer after 6 h palmitate treatment (D-I). ATP production was calculated as described in the Methods section. * = p < 0.05 Student’s t-test (A). In F-G, # = p < 0.05 vs. sh-NNT1; # = p < 0.05 vs. scramble. In I, * = p < 0.05 vs respective BSA group, $ = p < 0.05 vs. Palm-sh-NNT2, # = p < 0.05 vs BSA-Scramble, Two-way ANOVA + Tukey’ s post-test. OCR and ECAR plots are mean + SEM. Bar plots are mean + SD. Filled circles represent independent experiments.

An important regulator of mitochondrial NADP^+^/NADPH ratios is an enzyme located in the inner mitochondrial membrane, the nicotinamide nucleotide transhydrogenase (NNT), which promotes the reduction of NADP^+^ by the oxidation of NADH, powered by the mitochondrial inner membrane potential (ΔΨ). The NNT is, consequently, a node that integrates energy production (NADH pool and ΔΨ) and the mitochondrial antioxidant system (recycling of glutathione and thioredoxin) (Rydström, 2006). Furthermore, NNT has been shown to be inhibited by long chain acyl-CoAs, with palmitoyl-CoA as the most specific inhibitor (Rydström et al., 1971). We observed that palmitate treatment did not change NNT protein content after 6 hours (Fig 4B, Suppl Fig 5), although this does not exclude modulation of activity under these conditions.

Gameiro et al., 2013, described that NNT knockdown promotes a fuel switch from glutamine to glucose use by measuring their incorporation into TCA intermediates and lactate secretion. We speculated if palmitate-induced glycolytic activation could be related to NNT inhibition, and, thus, decreased NADPH content, and if NNT inhibition could exacerbate palmitate-induced glycolytic flux. To address this, we stably knocked down NNT using shRNA; the protein content was decreased to less than 50% of control levels (Fig 4B). Both clones tended to decrease basal OCRs, independently of palmitate (Fig 4C, p < 0.067, Two-way ANOVA). Interestingly, the proton-leak component of oxygen consumption is significantly reduced in the 28512 clone (i.e. NNT1, Fig 4D, 4F), which is compatible with NNT function, that allows for proton re-entry into the matrix. Non-mitochondrial respiration was increased in the 28541 clone (i.e. NNT2) compared to the control (Fig 4E, 4G). Furthermore, while there were no differences in total ATP production (Fig 4H), ATP from glycolysis is significantly increased in NNT2 KD (Fig 4I). This demonstrates that changing mitochondrial redox state either through palmitate treatment or NNT KD can promote glycolytic ATP production.

### 3.5. Oxidative disturbances promote glycolysis

Based on the results seen up to now, we hypothesized that redox signals from mitochondria could play a decisive role in the upregulation of ATP synthesis promoted by palmitate. We thus evaluated if the direct generation of mitochondrial oxidants could regulate glycolysis. To do so, we used the naphthoquinone analogue menadione, which increases superoxide formation and lowers NADPH levels (Smith et al., 1987). Menadione lead to increased OCRs as a consequence of enhanced proton leak (Fig 5A), a result which differs from the effects of palmitate and NNT KD seen previously. Menadione also enhanced glycolytic flow (Fig 5B), without changing total ATP production (Fig 5C). Interestingly, the contribution of glycolysis toward ATP production (Fig 5D) was almost 3-fold higher with menadione than in the DMSO control, suggesting that 6 hours of redox imbalance is enough to remodel cellular metabolism through glycolytic activation without compromising net ATP production.

**Figure 5:**
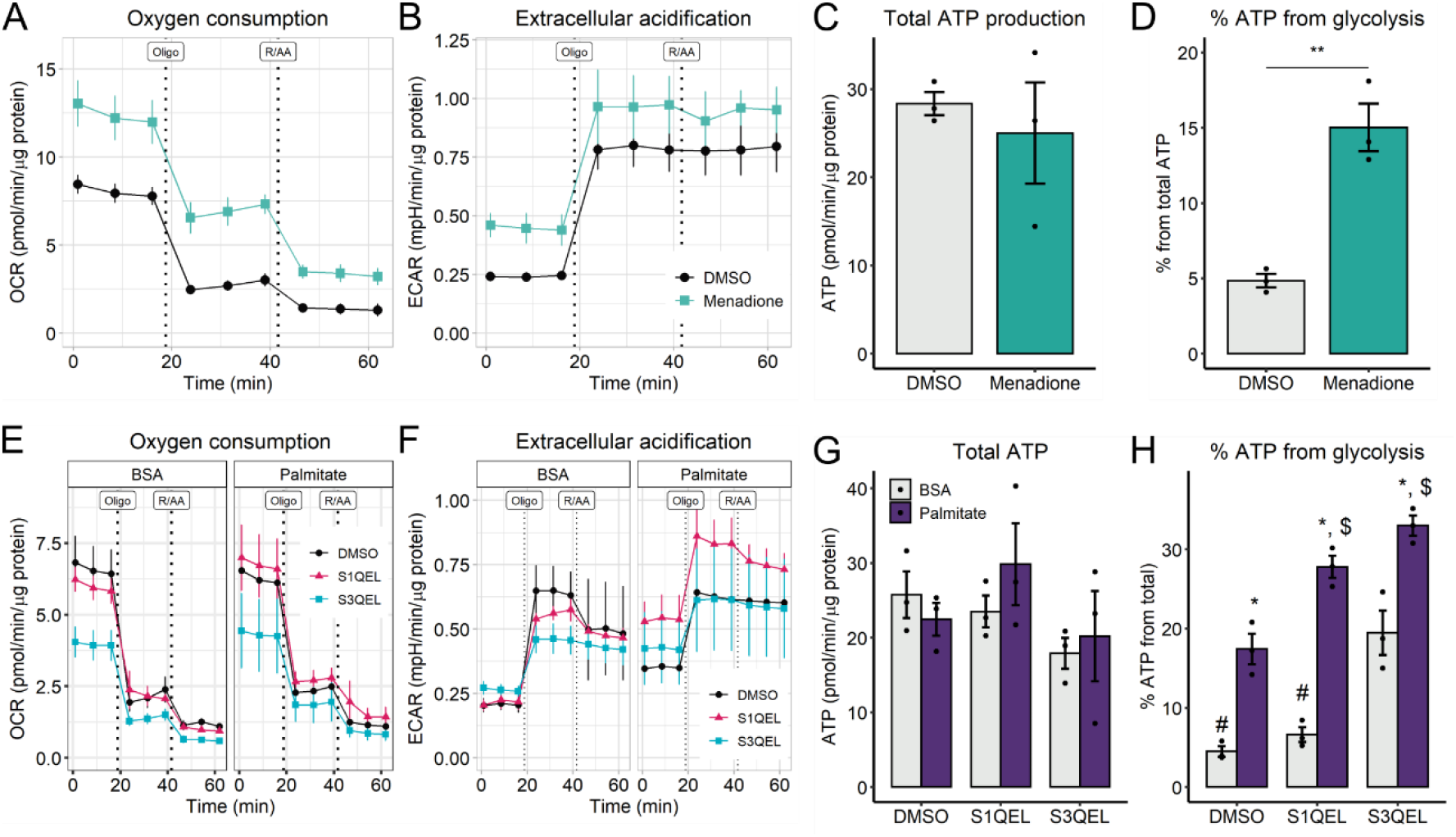
Site IQ of ETC inhibition exacerbated increased glycolytic flux promoted by palmitate. PLC/PRF/5 cells were incubated with palmitate/BSA (0.2 mM/ 0.25%) for 6 h and 10 μM menadione (A-D), 10 μM S1QEL, or 10 μM S3QEL (E-H), where indicated. Vehicle is 0.1% DMSO. Real-time OCRs and ECARs were measured in a Seahorse XFe24 analyzer after 6 h. ATP production was calculated as described in the Methods section. In C-D, p < 0.05 Student’s t test. In G-H, ** = p < 0.05 vs respective BSA; $ vs Palm-DMSO, # vs BSA-S3QEL, Two-way ANOVA + Tukey post-test. OCR and ECAR plots are mean + SEM. Bar plots are means + SD. Filled circles represent independent experiments.

To evaluate the contribution of mitochondrially-generated oxidants in palmitate-induced glycolysis, we co-incubated PLC cells with palmitate and S1QEL1.1 and S3QEL2 (suppressors of complex I and III site Q electron leaks, respectively). These are small molecules screened to block superoxide production from the electron transport chain without compromising electron flow to complex IV-bound oxygen (Brand et al., 2016; Orr et al., 2015). Despite this, in our cells, we observed that S3QEL, but not S1QEL (both at 10 μM), lowered oxygen consumption (Fig 5E), but did not affect total ATP production (Fig 5G). No significant effects were seen on ECARs either (Fig 5F). Interestingly, both modulators exacerbated glycolysis in palmitate-treated cells, and S3QEL also increased glycolytic flux in BSA controls (Fig 5H). These results suggest that both I_Q_ and III_Q_ oxidant production sites provide signals that control cytoplasmic ATP production, and I_Q_ seems to be a site sensitive to palmitate. Overall, these results demonstrate that palmitate and redox state are important determinants of glycolytic flux.

## 4. Discussion

Palmitate overload is well known to promote lipotoxicity in many cell types (Jheng et al., 2012; Kulkarni et al., 2016; Molina et al., 2009). Here, we aimed to understand the early metabolic effects of palmitate overload, and how these were related to mitochondrial morphological changes. We have previously found, in a BV2 microglial culture, that neither palmitate nor oleate overload for 24 hours promote changes in mitochondrial oxygen consumption. Additionally, significantly decreased glycolysis and distinct lipidic species accumulation was observed in this inflammatory cell line (Chausse et al., 2019).

Prior studies in hepatic cells (Bao et al., 2010; Egnatchik et al., 2014; Geng et al., 2020) suggested that palmitate promoted a late decrease in inner mitochondrial membrane potentials and increased oxygen consumption, secondary to calcium efflux from the ER and increased glutamine catabolism (Egnatchik et al., 2014). We should note, however, that while mitochondrial function in intact cells is reliably evaluated by oxygen consumption rates (Brand and Nicholls, 2011), uncalibrated mitochondrial inner membrane potential estimates are much more artifact-prone (Gerencser et al., 2012), particularly when overt changes in morphology occur (Kowaltowski et al., 2002, Kowaltowski et al., 2019). Our experiments here show that, surprisingly, within 1 hour of incubation, palmitate can strongly increase glycolytic flux. An associated increase in glycolytic ATP production occurred, as soundly confirmed by 2-deoxiglucose inhibition of ECARs, but without disturbing mitochondrial oxygen consumption for up to 6 hours (Fig 2). Under these conditions, added palmitate was not oxidized, since etomoxir, a CPT1 inhibitor which prevents uptake of the fatty acid into mitochondria (Divakaruni et al., 2018), had no effects on OCR or ECAR (Suppl Fig 2). However, cytosolic palmitate actions can rewire the TCA and promote higher usage of glutamine (Egnatchik et al., 2014), which may explain our results.

At 6 hours under the same conditions of palmitate treatment, mitochondria are predominantly fragmented (Fig 1). However, our substrate availability studies showed that fragmentation and the increase in glycolytic ATP production are dissociated: while palmitate-induced mitochondrial fragmentation required the presence of both glutamine and glucose, palmitate-induced increases in ATP production from glycolysis required only glucose (Fig 3). Glutamine starvation was described recently as a strong pro-fusion signal (Cai et al., 2018), so we speculate that glutamine and/or glucose starvation are hierarchically stronger signals that favor elongation (Gomes et al., 2011), compared to palmitate-induced fragmentation.

Carbon flux through glycerol-3-phosphate consumption may contribute toward the morphology-independent palmitate-induced increased glycolytic flux we observed. Indeed, glycerolipid synthesis has been recently shown to be increased by palmitate, resulting in diacylglycerol accumulation (Piccolis et al., 2019). This flux could explain why lactate production (Suppl Fig 1) was less stimulated by palmitate than glycolytic flux and ATP generation. Another reason for lower lactate production than expected by the glycolytic rates is that cells were in reductive imbalance compared to the BSA controls (Fig 4), with low NAD(P)H/NAD(P)^+^ ratios, while lactate formation is stimulated by NADH accumulation. Additionally, lactate could have been converted in other metabolites (e.g., pyruvate, alanine or glucose).

A possible node for the regulation of the metabolic switch promoted by palmitate is the interaction between adenosine monophosphate-activated protein kinase (AMPK) and CD36, which mediates palmitate entry into the cell. Palmitate interacts with CD36 and can palmitoylate it (Tao et al., 1996; Thorne et al., 2010; Zhao et al., 2018). This increases CD36 translocation to the plasma membrane and favors the formation of a complex with Fyn, Lyn and LKB1, preventing AMPK activation under chronic palmitate treatment conditions (Li et al., 2019; Zhao et al., 2018). Acute binding to palmitate, on the other hand, may alleviate LKB1 inhibition, activating AMPK and lipophagy (Li et al., 2019; Samovski and Abumrad, 2019; Samovski et al., 2015). Zhang and co-workers elegantly described the mild and early activation of AMPK upon glucose starvation before any changes in ATP levels, in a manner regulated by aldolase activity (a glycolytic enzyme). AMPK is fully activated when AMP/ATP ratios start to rise in combination with glutamine deprivation (Zhang et al., 2017). Curiously, cells deprived of glucose do not activate glycolysis upon palmitate after 6 hours, indicating that PLC cells may be unable to provide carbons to form glycolytic intermediates, even when total ATP production is not compromised (Fig 3). Interestingly, in more recent work, the group observed that AMPK is activated in a stratified manner in compartments, proportionally to the severity of nutrient stress (Zong et al., 2019).

Another clear metabolic effect of palmitate was an increase in total NAD(P) and a shift to a more oxidized redox state of the pyridine nucleotide pool. The ability of palmitate to both decrease NNT (observed by McCambridge et al., 2019) content and inhibit it (Rydström et al., 1971) make NNT a notable redox control node, integrating redox state and substrate use during lipotoxicity. Indeed, NNT is seminal in the control of NADPH content in the mitochondrial matrix, responsible for the maintenance of ~50% of the NADPH pool (Rydström, 2006). Since the discovery of a spontaneous NNT mutation in Jax C57BL/6J mice (Toye et al., 2005), NNT importance has been increasingly recognized. Mice lacking the functional enzyme are more sensitive to diet-induced metabolic diseases (Fisher-Wellman et al., 2016; Francisco et al., 2018; Navarro et al., 2017). In cell culture systems, NNT knockdown disrupts not only redox balance and bioenergetics (Yin et al., 2012), but also switches fuel preference from glutamine to glucose, by regulating the NADPH/NADP^+^ ratios, increasing glucose incorporation into the TCA cycle, lactate secretion, and sensitivity to glucose deprivation (Gameiro et al., 2013).

Indeed, when we constructed NNT KD cells, we found that palmitate treatment combined with NNT2 KD increased the participation of glycolytic ATP in total production (Fig 4). However, we also observed decreased basal OCRs, mainly due to decreased ATP production from the electron transport chain, which is not seen with palmitate treatment alone, but could reflect the importance of NNT in mitochondrial functional maintenance. Increased non-mitochondrial oxygen consumption also suggests an enhanced activity of cellular oxidases and/or the production of reactive oxygen species in the knockdown cells. Given these effects, it is unclear if NNT KD can directly induce glycolysis in our model. Ward et al., 2020, showed that the NNT KD did not alter redox balance in several non-small cell lung cancer cell lines, but lowered mitochondrial respiratory capacity, and diminished the ability to oxidize palmitate (Ward et al., 2020). This suggests that the disruptive effects of NNT may be context-dependent.

Despite the ambiguous role of the NNT, the significant change in NAD(P)^+^/NAD(P)H rations demonstrate that palmitate strongly impacts on redox state. Based on this, we investigated if glycolysis could be modulated by changes in mitochondrial oxidant production. Recently, Brand’s lab generated important tools to investigate the role of electron transport chain reactive oxygen species production in cell biology (Brand et al., 2016; Orr et al., 2015). They found small molecules, S1QEL and S3QEL, that block, specifically, the production of superoxide/hydrogen peroxide without disturbing electron flow to oxygen, nor the production of oxidants at other sites. We took advantage of these molecules and identified that palmitate-induced glycolysis is exacerbated by the inhibition of superoxide/hydrogen peroxide at site I_Q_ (Fig 5), a location within mitochondrial complex I that produces both superoxide and hydrogen peroxide toward the matrix. Site III_Q_ produces superoxide toward both the matrix and the intermembrane space. Curiously, S3QEL increased glycolysis in both BSA and palmitate groups. However, since OCRs were significantly inhibited by it, we cannot exclude that other effects may be associated with the increase. This respiratory inhibition was not observed with S1QEL, which nonetheless affected glycolytic ATP production (Fig 5). We can thus conclude that the production of oxidants in the I_Q_ site is associated with metabolic rewiring of glycolysis promoted by palmitate.

## 5. Conclusion

Our work uncovers important early bioenergetic effects promoted by palmitate overload. We found that palmitate relies on glutamine and glucose availability to promote mitochondrial fragmentation. However, mitochondrial ATP production is unperturbed by a high degree of mitochondrial network fragmentation and oxidative imbalance, while glycolysis is readily responsive to acute palmitate treatment, oxidative stress, and mitochondrial oxidant production. Interestingly, inhibition of superoxide/hydrogen peroxide production at complex I site I_Q_ was sensitive to palmitate and activated glycolysis. This indicates a novel, fatty-acid modulated, redox regulation of glycolytic ATP production.

## 6. Acknowledgments

The authors thank Camille Caldeira da Silva, Dr. Juan Pablo Muñoz, Dr. María Isabel Hernández-Alvarez, Jordi Seco, Dr. Sebastien Tosi, and Dr. Nikolaos Giakoumakis for excellent technical support and experimental help, and Dr. Bruno Chausse for manuscript revision. This work was funded by *Centro de Pesquisa, Inovação e Difusão de Processos Redox em Biomedicina* (CEPID Redoxoma) grant 2013/07937-8, FAPESP DD 2015/25862-0, MINECO (SAF2016-75246R), the Generalitat de Catalunya (Grant 2017SGR1015), CIBERDEM (“Instituto de Salud Carlos III”), the Fundación Ramon Areces (CIVP18A3942), the Fundación BBVA, Fundació Marató de TV3 (20132330), and CAPES, with no involvement of the funding agencies in the design of this study.

## 7. Author contributions

Conceived and designed the experiments: PK, AZ and AK. Performed the experiments: PK. Analyzed the data: PK. Contributed reagents/materials/analysis tools: AZ, AK. Wrote the manuscript: PK and AK. Reviewed final version of the manuscript: PK, AZ and AK.

