## Supplementary material for "Increased glycolysis is an early outcome of palmitate-mediated lipotoxicity"

#### Supplementary figure 1

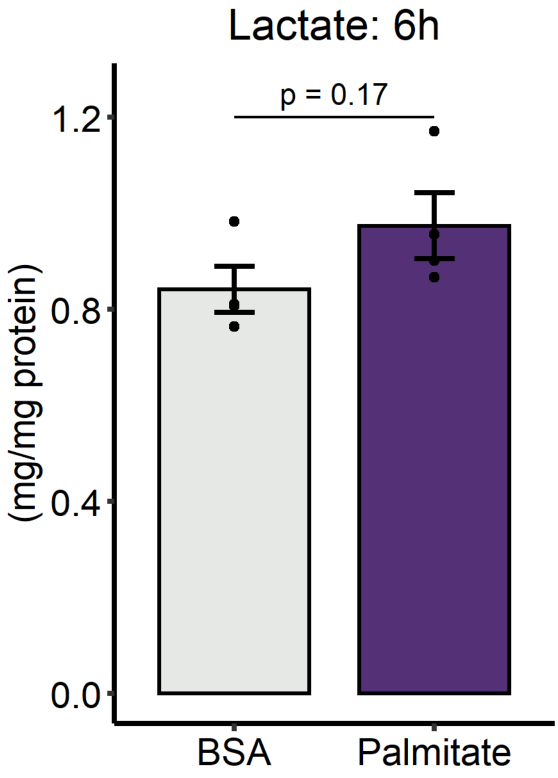


Suppl Fig 1. PLC/PRF/5 cells were incubated with palmitate/BSA (0.2 mM/ 0.25%) for 6 h and the supernatant was collected for lactate measurements. Student's t test. Data are mean + SD, and filled circles represent independent experiments.

Supplementary figure 2

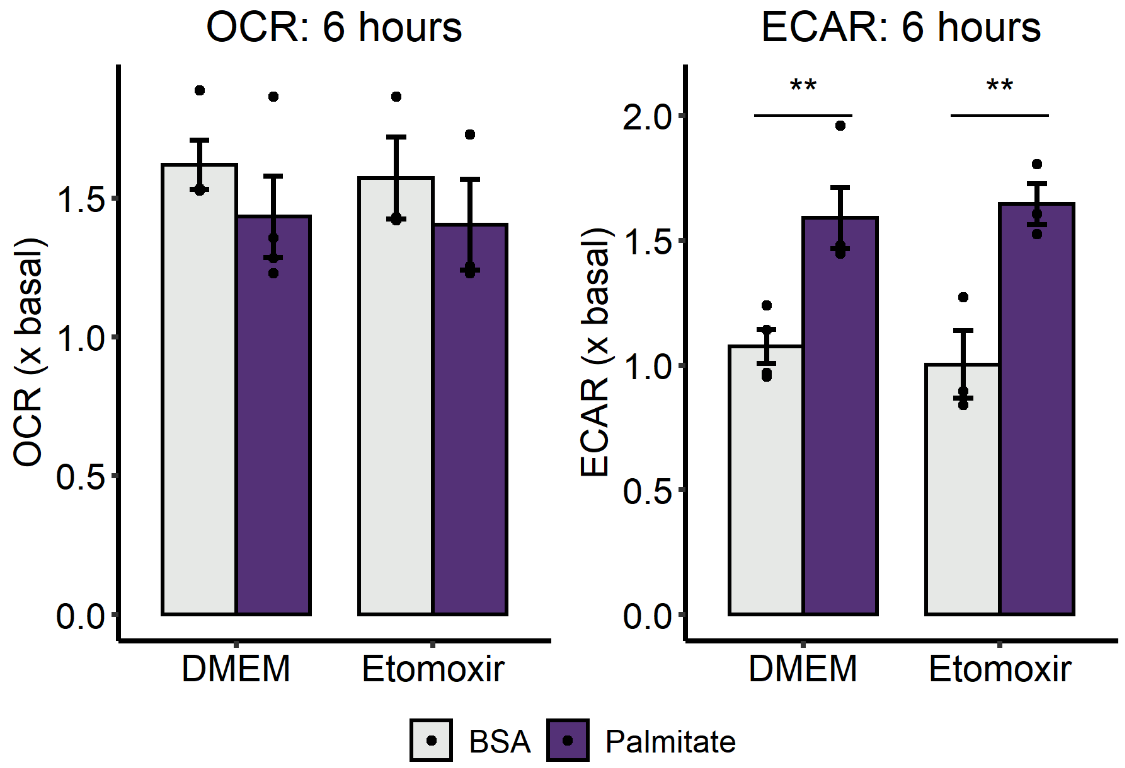


Suppl Fig 2. PLC/PRF/5 cells were incubated with palmitate/BSA (0.2 mM/ 0.25%) and 0.5 μM etomoxir for 6 h in a Seahorse XFe24. Basal OCRs and ECARs were measured and normalized by data pre-injection. Two-way ANOVA. $p < 0.01$ for palmitate effect. Data are mean + SD and filled circles represent independent experiments.

Supplementary figure 3

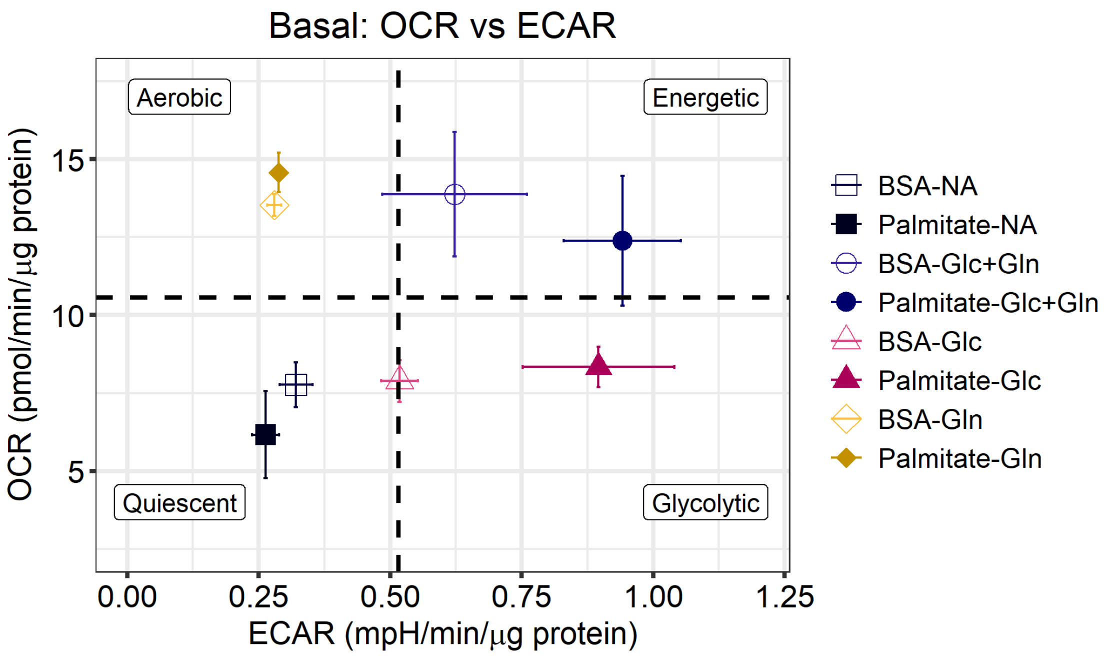


Suppl Fig 3. PLC/PRF/5 cells were incubated with palmitate/BSA (0.2 mM/ 0.25%) for 6 h in media without glucose nor glutamine (N/A), or in the presence of glucose and glutamine (Glc+Gln), or only glucose (Glc), or only glutamine (Gln). Basal OCR and ECAR were measured in Seahorse XFe24. Dashed lines represent all groups average. Data are mean + SEM (N = 3).

Supplementary figure 4

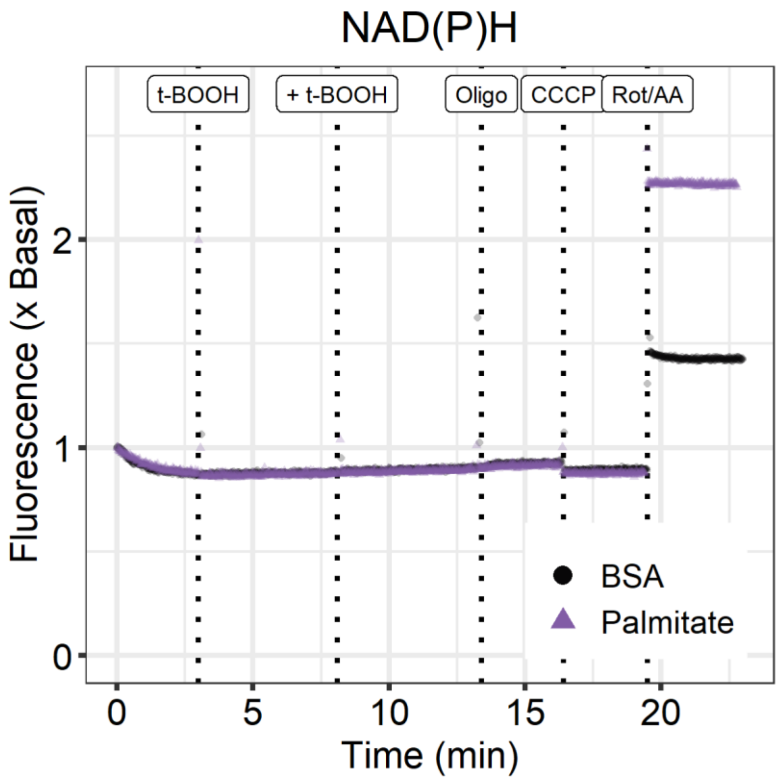


Suppl Fig 4. PLC/PRF/5 cells were incubated with palmitate/BSA (0.2 mM/ 0.25%) for 6 h. Representative experiment of intact cell NAD(P)H content measurement, modulated by the sequential injections of t-butyl hydroperoxide, oligomycin, CCCP, rotenone and antimycin A.

### Supplementary figure 5

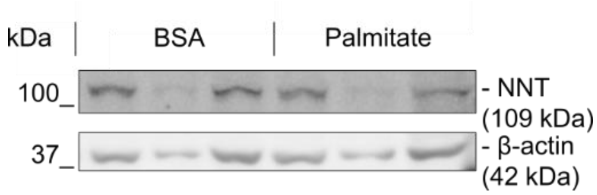


Suppl Fig 5. Suppl Fig 4. PLC/PRF/5 cells were incubated with palmitate/BSA (0.2 mM/ 0.25%) for 6 h and harvest for SDS-PAGE Western Blot. Representative bands for NNT.
